# PerfectlyAverage: a classical open-source software method to determine the optimal averaging parameters in laser scanning fluorescence microscopy

**DOI:** 10.1101/2025.03.17.643716

**Authors:** S. Foylan, L.M. Rooney, W. B. Amos, G.W. Gould, G. McConnell

**Affiliations:** Strathclyde Institute of Pharmacy and Biomedical Sciences, University of Strathclyde, 161 Cathedral Street, Glasgow, G4 0RE; MRC Laboratory of Molecular Biology, Cambridge Biomedical Campus, Francis Crick Avenue, Cambridge, CB2 0QH

**Author notes:** These authors contributed equally to this work (S. Foylan, L.M. Rooney).

**Keywords:** Laser scanning, microscopy, fluorescence, signal-to-noise, power spectrum, averaging

## Abstract

Laser scanning fluorescence microscopy (LSFM) is a widely used imaging method, but image quality is often degraded by noise. Averaging techniques can enhance the signal-to-noise ratio (SNR), but while this can improve image quality, excessive frame accumulation can introduce photobleaching and may lead to unnecessarily long acquisition times. A classical software method called PerfectlyAverage is presented to determine the optimal number of frames for averaging in LSFM using SNR, photobleaching, and power spectral density (PSD) measurements. By assessing temporal intensity variations across frames, PerfectlyAverage identifies the point where additional averaging ceases to provide significant noise reduction. Experiments with fluorescently stained tissue paper and fibroblast cells validated the approach, demonstrating that up to a 4-fold reduction in averaging time may be possible. Data are also presented to suggest that frame and line averaging with the same number of averages may produce different results, and this may influence the optimal number of averages. PerfectlyAverage is open source, compatible with any LSFM data obtained using line or frame averaging during image acquisition, and it is aimed at improving imaging workflows while reducing the reliance on subjective criteria for choosing the number of averages.

## 1. INTRODUCTION

Laser scanning fluorescence microscopy (LSFM) is a widely used technique in biological and materials sciences, offering high-resolution, multi-dimensional imaging capabilities^1^. However, LSFM images are inherently affected by noise sources such as photon shot noise and detector noise, which can degrade image quality. To address these challenges, averaging techniques are commonly employed to enhance the signal-to-noise ratio (SNR) and improve image clarity^2^.

Averaging in LSFM can be implemented through different approaches, with frame and line averaging being the most common methods^3^. Frame averaging involves capturing multiple frames of the same field of view and averaging the signal to reduce noise, while line averaging averages multiple signal readings along a scanned line. Both techniques are widely applied in confocal and multiphoton laser scanning microscopy to improve image SNR without increasing excitation power^4^.

The primary advantage of averaging is its ability to significantly enhance SNR by reducing random noise while preserving the underlying signal. In signal processing, the improvement in SNR is given by

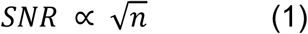

where 𝑛 is the number of lines or frames averaged, and in LSFM the SNR is given by

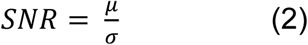

where 𝜇 is the mean intensity and 𝜎 is the standard deviation of the pixel values within the image^1^.

Averaging in LSFM is particularly beneficial for imaging weakly fluorescent specimens or specimens with low contrast, where noise can obscure fine structural details^5,6^. Despite its benefits and widespread use, averaging in laser scanning microscopy also has limitations. Increased acquisition time is a major drawback, as it can lead to specimen drift, image blurring, and motion artefacts, which are especially problematic in live cell imaging or time-sensitive experiments. Additionally, prolonged laser exposure can cause photobleaching in fluorescence microscopy and phototoxicity in living specimens^7^, potentially compromising their viability, and equation (1) does not take account of any photobleaching. These factors suggest that a careful balance between improving image quality and preserving specimen integrity is needed, yet in practice the number of frames used for averaging is mostly determined by subjective visual inspection of image quality.

Here, a classical software method termed PerfectlyAverage is presented to determine the optimal number of frames in laser scanning microscopy. By analysing temporal variations in pixel intensity across multiple frames, the method employs power spectral density (PSD) measurement to estimate the point at which additional frames no longer contribute significant new information and provides a recommended frame count for improved signal preservation and noise reduction. Power spectral density (PSD) analysis is particularly well-suited for evaluating image sequences in laser scanning microscopy because it provides a frequency-domain representation of noise and structural information^8^. Unlike time-domain analyses, which can be sensitive to local fluctuations, PSD enables a systematic assessment of noise characteristics by decomposing image information into spatial frequency components^8^. This allows for the identification of trends in noise reduction as more frames are averaged, helping to pinpoint the point at which additional frames cease to provide meaningful improvements. Furthermore, by incorporating spectral entropy derived from PSD, as has been previously applied in electroencephalography^9^, the method can objectively quantify structural consistency across frames, reducing reliance on subjective visual inspection. This makes PSD a powerful tool for determining the optimal number of frames for averaging, balancing noise reduction, and preserving image integrity while accounting for photobleaching effects. PerfectlyAverage also considers the measured SNR and is also resilient against the deleterious effects of photobleaching within user-defined limits, and it is open source.

## 2. METHOD

### 2.1 Specimen Preparation

Two specimens were prepared for LSFM imaging to assess the method. The first was a section of lens tissue paper stained with 10 µM Safranin O (S2255, Sigma-Aldrich) in ethanol mounted in a gelvatol medium, and the second was a fibroblast cell preparation. Fibroblast cells (3T3-L1, CL-173) were grown in vented capped tissue culture flasks containing DMEM (10567-014, Thermo Fisher) supplemented with 10% NBCS (26010066, Thermo Fisher) and 1% penicillin streptomycin (15140-122, Thermo Fisher) and 1% L-glutamine (21051040, Thermo Fisher) and were incubated at a temperature of 37°C in a 10% CO_2_ humidified cell incubator (Heracell VIOS CO_2_, Thermo Fisher). Cells were seeded at the desired density and incubated for 24 h at 37°C/10% CO_2_ to promote adherence and then rinsed twice with PBS (10010023, Thermo Fisher) prior to fixation in 4% paraformaldehyde (158127, Sigma-Aldrich) for 10 mins, and rinsing three times in PBS (10010023, Thermo Fisher). Cells were permeabilised using 500 µl of 0.1% Triton X-100 (T8787, Merck) in PBS at room temperature for 15 mins, then washed 3 times with PBS before adding two drops of ActinGreen 488 ReadyProbes to each coverslip (R37110, Thermo Fisher), and were incubated for 30 minutes at 37°C/10% CO_2_. The staining solution was removed, and the cells were washed three times in PBS (10010023, Thermo Fisher) before mounting in Prolong Glass AntiFade Mountant (P36980, Thermo Fisher). Both the stained lens tissue paper and the 3T3-L1 cell specimens were mounted between a microscope slide and a Type 1.5 coverslip and were allowed to set overnight at room temperature without ambient light before subsequent imaging.

### 2.2 Data Acquisition and Generation for software testing

An image dataset was generated in FIJI^10^ to simulate real data but with *a priori* knowledge of the noise characteristics. A 512 pixel x 512 pixel 16-bit image filled with noise was generated in FIJI. This image served as the first image in the stack to mimic an image with no averaging. This image was duplicated, and this second image was despeckled by applying a 3 pixel × 3 pixel median filter, and hence denoised, in FIJI. This served as the second image. This process was repeated to obtain progressively less noisy images with increasing image number, and these images were compiled into a single .tif stack.

Image data to evaluate the method were acquired using an upright microscope (DM6000CFS, Leica) coupled to a laser scanning system (SP5, Leica) controlled by software (LAS-AF v2.7.7.12402, Leica). Image stacks of each specimen were acquired with increasing number of line and frame averaging, starting with a minimum of no line or frame averaging up to 1024 lines and up to 64 frames in a geometric sequence. These values were the maximum numbers of line and frame averages possible with the image acquisition software. All data were acquired at a scan speed of 400 Hz and were saved in the proprietary format (.lif): these were converted to individual .tif files and compiled into .tif stacks using FIJI. These LSFM data served as the input to the software code.

For imaging of the tissue paper stained with Safranin O, a 488 nm laser was used for excitation of fluorescence, which was detected between 500-550 nm with a spectral detector. A 10x/0.4 numerical aperture dry objective lens (10x/0.4 HC PL APO, Leixa) was used for imaging. The laser power at the specimen plane was measured with a power meter (Nova II, Ophir Photonics) to be 0.7 mW, and the detector gain was set to 900. Images of 4096 pixels x 4096 pixels were captured with a pixel size of 360 nm to satisfy the Nyquist-Shannon sampling criterion^11^.

For imaging of the 3T3-L1 cells prepared with ActinGreen dye, the same 488 nm laser was used for excitation of fluorescence, which was also detected between 500-550 nm with a spectral detector. A 20x/0.7 numerical aperture dry objective lens (20x/0.7 HCX PL APO CS, Leica) was used with a digital zoom of 2 for imaging. The laser power at the specimen plane was measured to be 0.2 mW, and the detector gain was set to 1050. Images of 1024 pixels x 1024 pixels were captured at a rate of 400 Hz and with a pixel size of 360 nm, and the Nyquist-Shannon sampling criterion was not satisfied^11^. The laser power at the specimen plane was set to a deliberately low value and the detector gain set to a high value to provide a poor SNR at low numbers of frame or line averaging to best challenge the PerfectlyAverage software.

### 2.3 Signal-to-Noise Ratio (SNR) Calculation

The signal-to-noise ratio (SNR) was computed for each image in the chosen .tif stack using the equation (2). A zero-noise safeguard was implemented to prevent division errors, ensuring numerical stability when processing images with minimal intensity variation.

### 2.4 Photobleaching Correction

To correct for photobleaching, the mean intensity of each frame was calculated. Photobleaching was considered as an exponential decay process^12–14^, with an exponential decay function fitted to the mean intensity values across the image sequence:

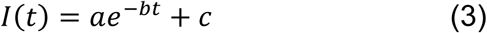

where 𝐼(𝑡) represents the mean intensity at frame 𝑡, and 𝑎, 𝑏, and 𝑐 are fitting parameters. The curve-fitting procedure was performed using the Levenberg-Marquardt algorithm^15^ as implemented in SciPy’s *curve_fit* function. Each frame was subsequently corrected by applying a normalisation factor derived from the fitted decay function, ensuring the corrected intensities remained proportional to their original values.

### 2.5 Power Spectrum Domain Analysis

A PSD analysis was conducted to examine frequency components within the image frames. In Python, the two-dimensional fast Fourier transform (FFT) was applied to each image, followed by a shift of the zero-frequency component to the centre^16^. The power spectrum was computed as:

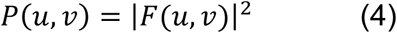

where 𝐹(𝑢, 𝑣) represents the Fourier-transformed image. The spectral entropy was then calculated as a measure of frequency distribution randomness. The PSD values were normalised to create a probability distribution, and the Shannon entropy was calculated^17^:

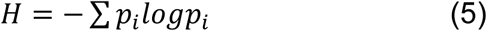

where 𝑝_i_ denotes the normalised power spectral density values.

Spectral entropy quantifies the complexity and randomness of frequency distributions within an image, providing insight into texture and structural variations^17^. A peak stability detection algorithm was employed to determine the optimal frame based on spectral entropy variation over time.

### 2.6 Optimal Frame Selection

To identify the optimal frame in the chosen dataset, both SNR and frequency-domain stability measurements were performed. The optimal frame based on SNR was selected by identifying the maximum SNR within the frames that met the user-defined photobleaching threshold. The optimal averaging conditions were identified by determining the point at which the normalised spectral entropy stabilised, using a moving average smoothing function with a window size of five frames and a stability threshold of 0.1. The choice of threshold in measuring PSD significantly influences the selection of the optimal frame for averaging based on PSD analysis. A lower threshold (e.g., 0.01) results in a readout of the earliest point where the rate of change in spectral entropy is minimal. While this approach is sensitive, it may prematurely identify an optimal frame before true stabilisation has occurred. Increasing the threshold (e.g., to 0.1) allows for greater tolerance to minor fluctuations in spectral entropy, thereby ensuring that the selection of the optimal frame aligns with a more sustained plateau in PSD values. This adjustment reduces the risk of prematurely selecting a frame where noise reduction is still improving, leading to a more balanced trade-off between averaging and noise suppression. Frames exhibiting the highest SNR were prioritised to ensure maximal signal clarity, while frequency stability was used to confirm minimal structural fluctuations. The integration of these two criteria allowed for a comprehensive selection of the ideal frame in the imaging sequence.

### 2.7 Software and Data Presentation

All code was developed using Python (v3.12) within the Spyder (v5.5.1) integrated development environment, managed via Anaconda Navigator (v2.6.3). The software was run on a Dell XPS 13 9370 laptop computer with an i7-8550U 1.8 GHz processor and 16 GB RAM, with a Microsoft Windows 10 Pro operating system. A Python version and a standalone Windows executable of PerfectlyAverage were produced.

Upon running the software, a dialog box asks the user to choose the dataset to be analysed. This is advised to be a dataset from the same or a similar specimen to be studied, using the preferred LSFM in an adjacent region of the preparation. Images are acquired with an increasing number of averages and data are saved as described above. Upon execution of the software code, the user is prompted to specify an acceptable level of photobleaching, expressed as a percentage decrease in signal intensity. This input ensured that the analysis focused on frames retaining sufficient fluorescence intensity for accurate interpretation. The photobleaching threshold was set to exclude frames where the intensity dropped below a user-defined percentage of the initial value, thereby preventing overcorrection or inclusion of frames with excessive signal degradation.

Upon execution, results were visualised using Matplotlib^18^, displaying normalised mean intensity, SNR, and spectral entropy using PSD across frames. The optimal frames determined from SNR and frequency-domain analyses were highlighted, with a summary of the optimal frame selection criteria and photobleaching correction parameters presented to the user. This interface was designed to provide a user-friendly mechanism for interpreting the results.

To produce the cropped region of datasets for analysis of the dependence of the image size on the number of averages recommended by PerfectlyAverage, a custom Python script was produced to automate this process.

## 3. RESULTS

An example image dataset and the output of the PerfectlyAverage software code is shown in Figure 1. Since these input data are strictly not averaged, the x-axis has been updated accordingly from the conventional PerfectlyAverage output to indicate despeckling rather than averaging. The simulated data comprised a 512 pixel x 512 pixel 16-bit image is shown in Figure 1(A), with progressive despeckling and hence denoising in subsequent frames shown in Figures 1(B-L). These data are shown with the ‘Glasbey’ false-colour look up table for enhanced contrast^19^. These images were compiled as an image stack and used with the Python version of PerfectlyAverage. An acceptable limit of 10% photobleaching was chosen, and the plot of the mean intensity, SNR and PSD are shown in Figure 1(M). The mean total processing time to run the code with these data was under 0.3 seconds on a standard laptop computer. The time to load the file was 0.008 ± 0.002 seconds, the photobleaching correction execution time was 0.026 ± 0.007 seconds, the time taken to calculate the SNR was 0.038 ± 0.010 seconds, and the PSD calculation took 0.199 ± 0.051 seconds.

**Figure 1.**
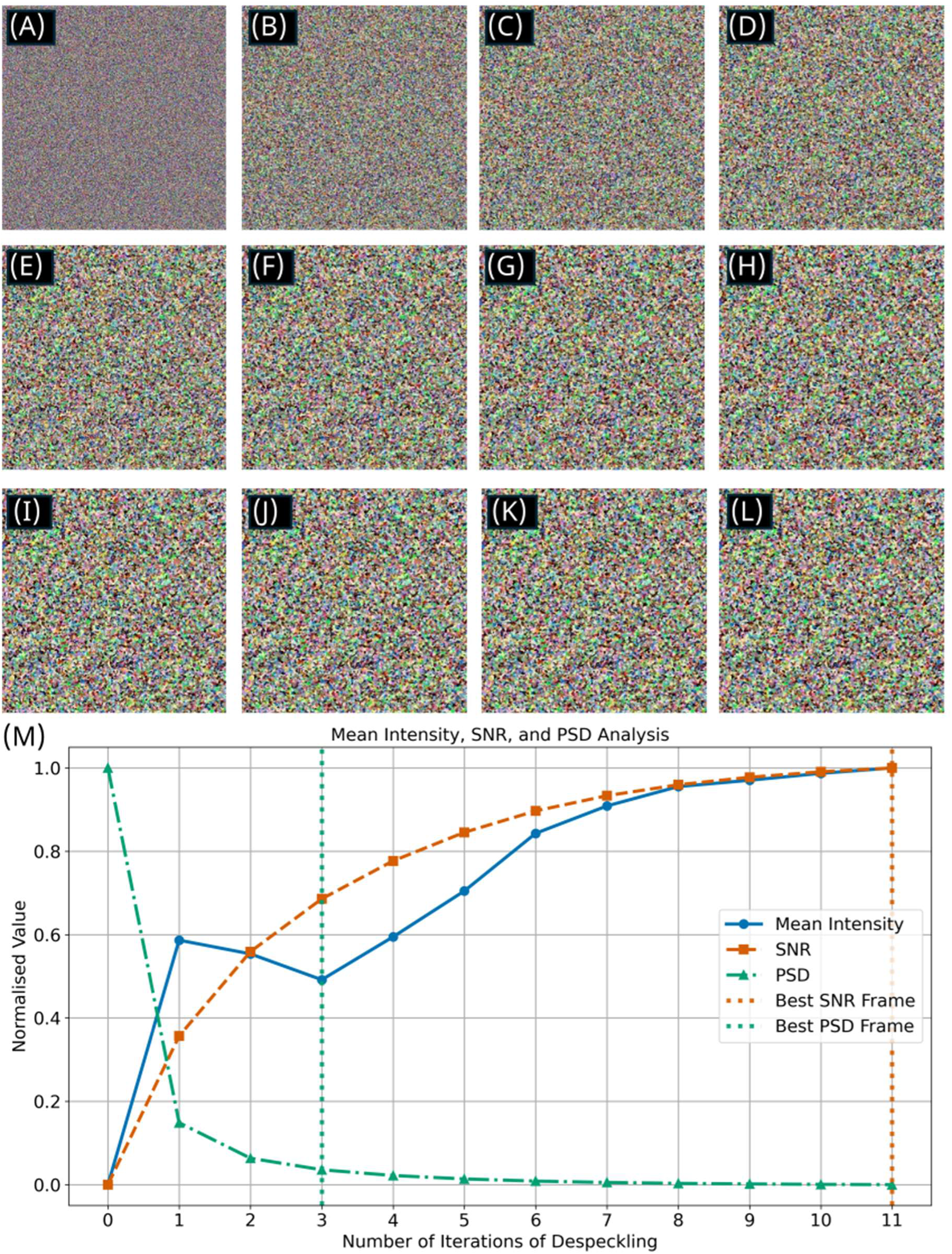
(A) Image generated in FIJI, comprising 512 pixels x 512 pixels and filled with noise, displayed using ‘glasbey’ false colour look up table. (B)-(L) show progressive despeckling of (A) performed in FIJI, with a single application of despeckling performed on each successive image. Images (A)-(L), compiled as a .tif stack, served as the input stack to the PerfectlyAverage software code. (M) Output plot from the PerfectlyAverage software code based on input images (A)-(L), for a chosen photobleaching limit of 10% (x-axis amended to accurately describe despeckling of simulated data rather than averaging of image data). Based on the signal-to-noise (SNR) ratio with respect to this photobleaching limit set by the user, the optimal parameter is defined as the input image with 11 iterations of despeckling, shown in 2(L), but power spectral density (PSD) analysis reveals that the optimal output image is that obtained with only three iterations of despeckling, shown in 2(D).

As expected for this simulated dataset, the SNR improves with the amount of despeckling applied. While this is not a simulated fluorescence image and technically has not been averaged in the conventional sense it has been progressively denoised and the mean intensity of the dataset varies nonlinearly with the amount of despeckling applied. With a photobleaching limit of 10%, which in this instance can be considered as an equivalent decrease in image signal amplitude of 10%, the optimal result is obtained with the final image in the stack. Since there is no photobleaching – indeed, the mean intensity increases at higher values – and the SNR increases nonlinearly, the photobleaching is not a limiting factor in this analysis. However, consideration of the PSD suggests that fewer denoising steps could be used to achieve a good imaging result and minimising exposure to light in practical imaging. In this example, the PSD reaches a plateau after 3 iterations of despeckling, suggesting that less averaging could be considered optimal for denoising.

Figure 2 shows the input LSFM images of lens tissue paper stained with Safranin O, and the output of the PerfectlyAverage code used with 10% photobleaching limit. Figure 2(A) shows the full field of view, with a white box highlighting a region of interest (ROI). All images are shown with the ‘16 colors’ look up table. This ROI is digitally zoomed and shown in images Figures 2(B)-(H). Figure 2(B) shows the ROI with no frame averaging, and images (C)-(H) show increasing numbers of averaged frames up to 2^6^=64 images. The scale bar in image 2(A) is 200 µm, and the scale bar for images 2(B)-(H) is 20 µm. Through subjective visual inspection, as expected image 2(B) has the most noise, and improvements in SNR may still be occurring at higher image numbers. The output of the PerfectlyAverage code, shown in Figure 2(I), confirms this observation, but the mean intensity responds nonlinearly, with some photobleaching. Based on the user-defined photobleaching limit of 10% and considering the SNR the ideal number of averaged images is 2^1^=2, corresponding to image 2(C). However, by analysis of the PSD, the ideal number of frame averages is 2^4^=16.

**Figure 2.**
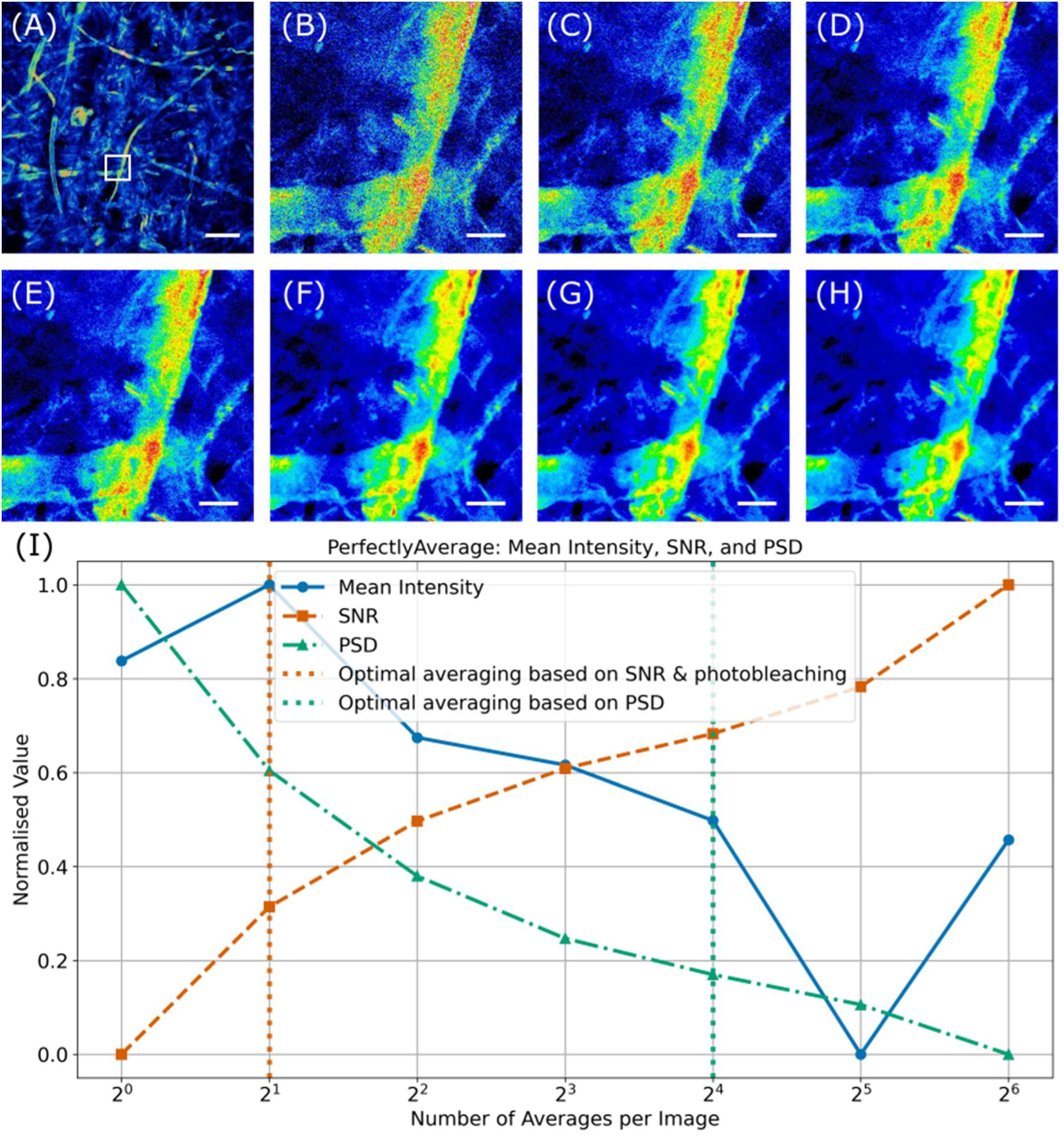
(A) Laser scanning fluorescence microscope image of lens tissue paper stained with Safranin O, shown using the ‘16 colors’ false colour look up table. A white box highlights a region of interest (ROI), scale bar = 200 µm. (B)-(L) show digital zooms of this ROI with (B) no frame averaging, and (C)-(H) showing increasing frame averaging up to 2^6^=64 frames, scale bars = 20 µm. The full images (and not the ROIs) served as the input to the PerfectlyAverage algorithm. (I) Output plot from the PerfectlyAverage software code based on input images for a chosen photobleaching limit of 10%. Based on the signal-to-noise (SNR) ratio with respect to this photobleaching limit set by the user, the optimal image is defined as the input image with an average of 2^1^=2, shown in 2(C), but power spectral density (PSD) analysis revealed that the optimal output image was obtained with frame averaging of 2^4^=16 frames per image, shown in 2(F).

Supplementary Figure 1 shows a plot of the optimal number of averages per image based on SNR and photobleaching, shown in magenta, and the optimal number of averages per image based on the PSD for the same frame averaged dataset presented in Figure 2, but with a central cropped region ranging from 8 pixels in diameter to no cropping, taking into account the full 4096 x 4096 pixel image size during the measurement. As the diameter of the image decreases below 512 pixels, the optimal number of averages when considering SNR and photobleaching increases rapidly from 2 images to 64 images, whereas the optimal number of averages remains constant at 16 irrespective of the image diameter. These data suggest that it may be possible to acquire data from sub-regions of the field of view, e.g. using ROI scanning, for use with PerfectlyAverage and PSD measurement for robust and rapid assessment of the optimal number of averages in practical imaging studies.

Figure 3 shows results of line averaging of the same specimen, again with a 10% photobleaching limit defined by the user. Figure 3(A) shows the full field of view, with a white box highlighting a region of interest (ROI). This ROI is digitally zoomed and shown in images Figures 3(B)-(H). All images are shown with the ‘16 colors’ look up table. Figure 3(B) shows the ROI with no line averaging, and images (C)-(H) show increasing numbers of line averaging per frame up to 2^6^=64 images. Visually, the images in Figures 2 and 3 appear similar, with decreasing noise at higher frame or line averages, as expected. However, with the same user-defined photobleaching limit applied to the line averaged dataset, with reference to the SNR the ideal number of line averages per image is 2^5^=32, which is a 16-fold increase on the same parameter measured for the specimen imaged with frame averaging. The ideal number of line averages determined by the PSD, however, is 2^4^=16, which is the same value determined by the PerfectlyAverage software code for frame averaging, as shown in Figure 2(I).

**Figure 3.**
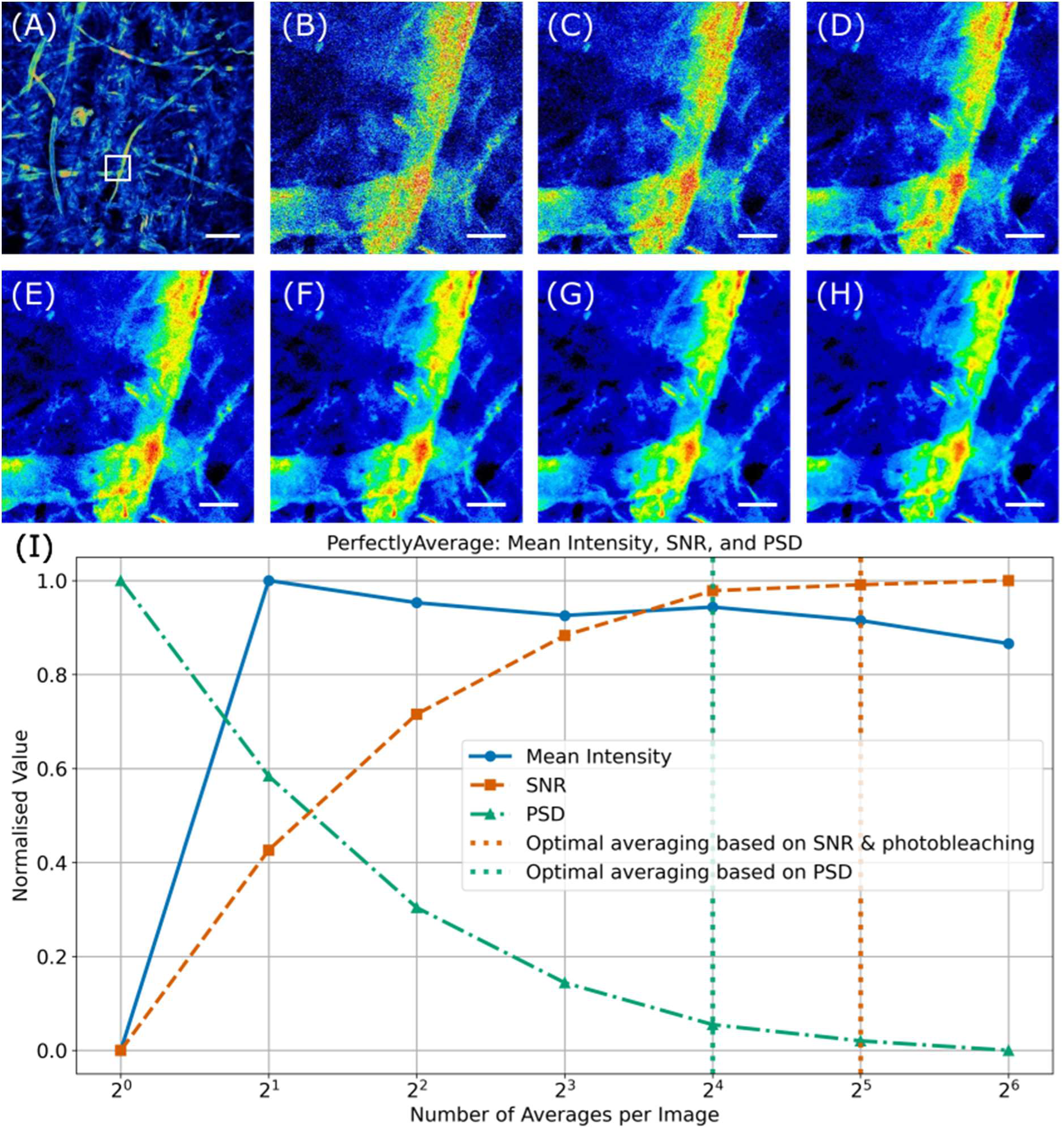
(A) Laser scanning fluorescence microscope image of lens tissue paper stained with Safranin O, shown using the ‘16 colors’ false colour look up table. A white box highlights a region of interest (ROI), scale bar = 200 µm. (B)-(L) show digital zooms of this ROI with (B) no line averaging, and (C)-(H) showing increasing line averaging up to 2^6^=64 frames, scale bars = 20 µm. The full images (and not the ROIs) served as the input to the PerfectlyAverage algorithm. (I) Output plot from the PerfectlyAverage software code based on input images for a chosen photobleaching limit of 10%. Based on the signal-to-noise (SNR) ratio with respect to this photobleaching limit set by the user, the optimal image is defined as the input image with an average of 2^5^=32, but power spectral density (PSD) analysis revealed that the optimal output image was obtained with frame averaging of 2^4^=16 frames per image. The optimal number of line averages (16 lines) obtained using PSD for line averaging matched the optimal number of frame averages (16 frames), as shown in Figure 2.

Figure 4 shows results of line averaging of the fixed 3T3-L1 cell specimen stained with ActinGreen, with a user-defined 5% photobleaching limit. Figure 4(A) shows the full field of view, with a white box highlighting a region of interest (ROI). This ROI is digitally zoomed and shown in images Figures 4(B)-(P). Figure 4(B) shows the ROI with no line averaging, and Figures 4(C)-(P) show increasing numbers of line averaging per frame up to 2^10^=1024 averaged lines per image. All images are shown with the ‘Fire’ look up table [FIJI]. Based on the user-defined photobleaching limit of 5% and considering the SNR the ideal number of averaged images is 2^5^=32, corresponding to Figure 4(k). However, analysis of the PSD suggests that the ideal number of line averaging is 2^3^=8, shown here in Figure 4(E). Reducing the number of line averages from 2^5^ to 2^3^ reduces the overall imaging time by a factor of 2^2^: given the scan speed of 400 Hz and the image size of 1024 pixels x 1024 pixels for these data a total image capture time of 82 seconds for 2^5^ images based on the SNR and photobleaching limit would reduce to 20.5 seconds for 2^3^ images by consideration of the PSD.

**Figure 4.**
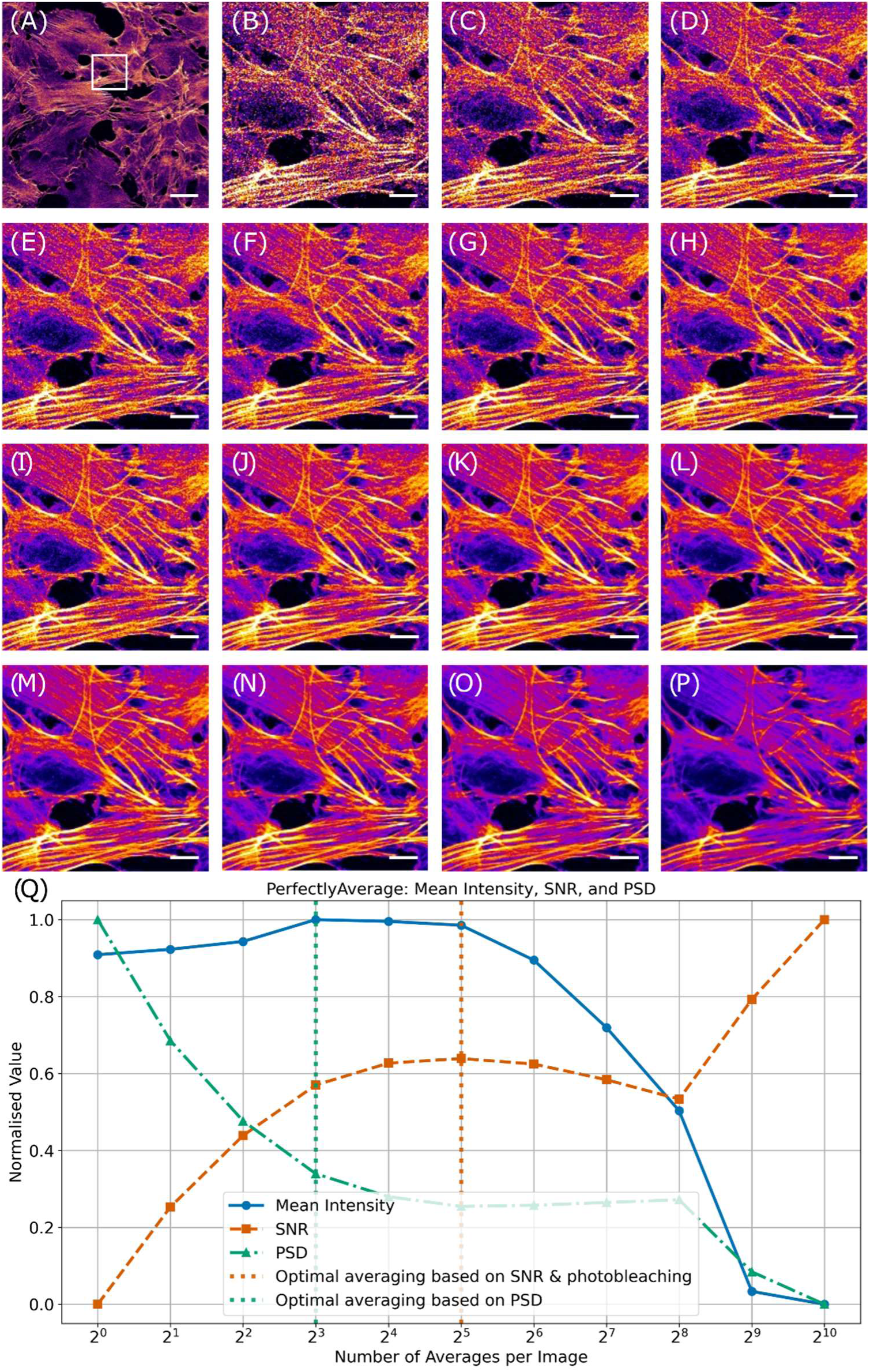
(A) Laser scanning fluorescence microscope image of fixed 3T3-L1 cells stained with ActinGreen, shown using the ‘Fire’ false colour look up table. A white box highlights a region of interest (ROI), scale bar = 50 µm. (B)-(P) show digital zooms of this ROI with (B) no line averaging, and (C)-(P) showing increasing line averaging up to 2^15^=32,768 frames, scale bars = 10 µm. The full images (and not the ROIs) served as the input to the PerfectlyAverage algorithm. (I) Output plot from the PerfectlyAverage software code based on input images for a chosen photobleaching limit of 5%. Based on the signal-to-noise (SNR) ratio with respect to this photobleaching limit set by the user, the optimal image is defined as the input image with an average of 2^5^=32, but power spectral density (PSD) analysis revealed that the optimal output image was obtained with frame averaging of 2^3^=8 frames per image.

## 4. DISCUSSION

PerfectlyAverage presents a systematic software approach for determining the optimal number of frames for averaging in LSFM. Using SNR measurements and PSD analysis, this method identifies the point at which additional frame averaging ceases to provide significant noise reduction, while also considering the effects of photobleaching. This is likely to have the greatest impact for LSFM images comprised of a large number of pixels, such as Mesolens data where LSFM images of up to 24000 pixels x 24000 pixels are produced^20^, and in other gigapixel-scale LSFM methods for imaging at the mesoscale^21,22^.

The application of the PerfectlyAverage algorithm suggests that relying solely on SNR for optimising line or frame averaging may lead to excessive image acquisition, resulting in unnecessary photobleaching and increased imaging times. Instead, PSD analysis provides a more robust criterion for determining the optimal frame count, often requiring significantly fewer averages than SNR-based selection. The results from both simulated and experimental datasets indicate that frame and line averaging can yield different optimal numbers, further emphasising the need for a more objective and systematic approach to determining averaging parameters in LSFM image acquisition.

The primary implication of these findings is that traditional approaches to determining frame averaging in LSFM, which typically rely on subjective visual inspection or SNR maximisation, may be sub-optimal. The use of PSD analysis offers an alternative, mathematically-grounded method for balancing noise reduction and photobleaching. PSD analysis provides insight into the frequency-domain characteristics of image noise and structural consistency, revealing when further averaging ceases to add valuable image information. The ability to establish an objective, reproducible criterion for optimal frame averaging represents a step forward in designing experiments, standardising LSFM imaging workflows, and communicating methods and results^23–26^.

Additionally, these results demonstrate that frame averaging and line averaging can produce different noise-reduction dynamics. The discrepancy between these two approaches suggests that the spatial characteristics of noise in LSFM images influence the effectiveness of averaging techniques. This observation has important implications for microscopy users, as it highlights the need to carefully consider whether frame or line averaging is more appropriate for a given imaging study.

The findings of this study align with prior research on noise reduction in LSFM imaging, which has largely focused on the trade-off between SNR improvements and photobleaching effects^1,6^. These previous studies have shown that averaging improves SNR proportionally to the square root of the number of frames averaged, but PerfectlyAverage adds value by considering user-defined photobleaching limits and frequency-domain analysis to optimise frame selection.

The use of PSD-based noise assessment has been applied in other fields, such as magnetic resonance imaging^27^ and atom probe tomography^28^, but its application to LSFM has been limited. Some prior works have suggested alternative methods for reducing noise, such as deep learning-based denoising techniques^29^, but these approaches often require extensive training datasets from many different specimens and specific computational resources may be needed. In contrast, PerfectlyAverage provides an easily implementable, open-source solution that can be readily adopted by LSFM users without requiring machine learning expertise.

Additionally, the observation that line and frame averaging yield different results adds a new perspective on data acquisition in LSFM. Previous studies have largely treated these two techniques as interchangeable^1^, but the data obtained using the PerfectlyAverage method suggests that line averaging may produce a different degree of noise suppression than frame averaging. This may arise from variations in noise autocorrelation across different spatial scales, and this could be the subject of further study.

Limitations of PerfectlyAverage must be acknowledged. Firstly, the implementation of PerfectlyAverage assumes a stationary noise process, which may not be universally applicable across all LSFM imaging conditions^30^. Some biological samples may exhibit dynamic fluorescence variations due to photophysical effects such as reversible photoswitching^31^ that are not fully captured by PerfectlyAverage. Additionally, the study focuses primarily on fluorescence-based LSFM imaging; the applicability of PerfectlyAverage to other image contrast modalities, such as scanned transmission or reflectance^32^, where photobleaching is unlikely to feature, remains to be tested.

Another limitation is that the selection of the photobleaching threshold in PerfectlyAverage is user-defined. While this allows for flexibility, it introduces a degree of subjectivity into the process. Future refinements to the PerfectlyAverage algorithm could incorporate automated photobleaching estimation, potentially improving the reproducibility of results across different experimental conditions.

PerfectlyAverage also requires an input dataset to estimate the optimal averaging conditions. This will be both microscope and specimen dependent. As such, it is recommended that in practice the user obtains frame and/or line averaged data for the purpose of applying PerfectlyAverage at a region of the specimen different to those which will form the area for routine study. As the data here have shown, different specimen types will yield different recommendations for averaging, and PerfectlyAverage should be applied at the outset of any study where averaging is needed, but Supplementary Figure 1 suggests that it may be possible to reduce acquisition time of these data by reducing the pixel number of the image dataset. Future research could explore whether machine learning approaches could complement PerfectlyAverage by dynamically adjusting PSD thresholds based on specimen properties or microscope configuration.

## 5. CONCLUSIONS

The development of PerfectlyAverage addresses a long-standing challenge in LSFM imaging by providing an objective, reproducible method for optimising frame and line averaging conditions for routine study. By integrating PSD analysis, SNR calculations, and photobleaching corrections, this method helps to determine the point at which averaging ceases to provide benefit and it enhances experimental efficiency by reducing unnecessary frame acquisition.

By offering an open-source software solution that is compatible with any LSFM, the aim is to make this tool widely accessible to the microscopy community to encourage adoption, integration into existing tools, further development to extend capability, and to overcome the current reliance on subjective or biased methods.

## Acknowledgements

This work was supported by the Medical Research Council (MR/K015583/1), the Biotechnology and Biological Sciences Research Council (BB/X005178/1), the Engineering and Physical Sciences Research Council (EP/I006826/1), and the Leverhulme Trust.

## Data availability statement

All software and the data that supports the findings of this study are openly available at the University of Strathclyde KnowledgeBase: https://doi.org/10.15129/919393bf-7735-4f47-90cd-ab09ddb0e8a9.

---

**Supplementary Figure 1.**
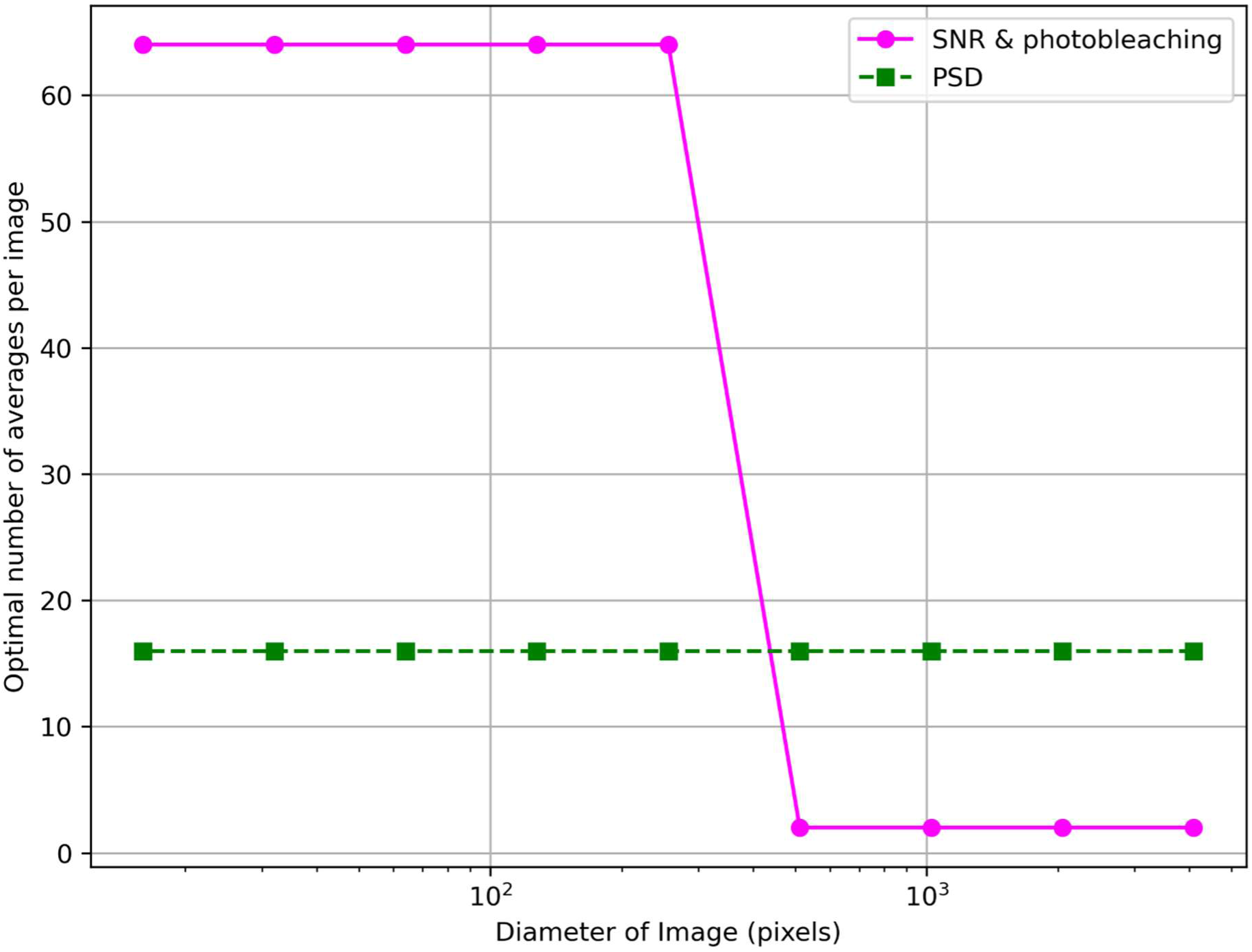
Plot of the optimal number of averages per image based on signal-to-noise (SNR) and photobleaching, shown in magenta, and the optimal number of averages per image based on the power spectral density (PSD) for the same frame averaged dataset presented in Figure 2, but with a central cropped region ranging from 8 pixels in diameter to no cropping, measuring the full 4096 x 4096 pixel image. As the diameter of the image decreases below 512 pixels, the optimal number of averages when considering SNR and photobleaching increases rapidly from 2 images to 64 images, whereas the optimal number of averages based on PSD measurement remains constant at 16, irrespective of the diameter of the image.

